# MF-ProtDisMap: protein real-valued distance prediction with fusion of sequence and coevolutionary features

**DOI:** 10.1101/2025.06.17.660102

**Authors:** Yufei Zhang, Suyang Zhong, Shenghui Xu, Zhumei Wang, Chengzhi Xin, Fei Ni, FangFang Yan, Xu Lu, Silong Sun, Hongwei Wang, Liang Zhang

**Affiliations:** College of Information Science and Engineering, Shandong Agricultural University, Tai’an, Shandong, China; State Key Laboratory of Wheat Improvement, College of Agronomy, Shandong Agricultural University, Tai’an, Shandong, China; Pen-Tung Sah Institute of Micro-Nano Science and Technology, Xiamen University, Xiamen, China; State Key Laboratory of General Artificial Intelligence, BIGAI Beijing, China

**Author notes:** Correspondence to (L.Z.); (H.W.). These authors contributed equally to this work (Yufei Zhang, Suyang Zhong).

**Keywords:** protein structure prediction, real-valued distance, protein language model, feature representation, diffusion model

## Abstract

The precise estimation of protein inter-residue distances is essential for high-accuracy protein structure modeling. Currently, the prediction methods are predominantly based on MSA-derived coevolutionary features or language model-based sequence features. To effectively leverage the strengths of both methods, this study developed MF-ProtDisMap (Multi-Feature Protein Distance Map), an integrative framework that effectively combines both feature types to achieve superior real-valued distance prediction. Briefly, MSA Transformer is employed to extract the coevolutionary features from protein multiple sequence alignments, whereas ESM2 is used to capture long-range interactions and sequence-level features. To reduce the computational cost while maximumly represent fused feature information, we implement group pooling for feature dimensionality reduction and introduce Diff-former—a novel module combining a diffusion model with a triangular attention mechanism to enhance representation learning. MF-ProtDisMap achieved a MAE of 2.20 Å and an RMSE of 3.40 Å in the protein real-valued distance prediction task. The predicted distances can be converted into contact results, achieving ROC and PR values of 84.56% and 81.01%, respectively. These results demonstrate that MF-ProtDisMap outperforms state-of-the-art real-valued protein distance methods.

## 1. Introduction

Recent breakthroughs in deep learning have greatly advanced the field of protein structure prediction, enabling unprecedented accuracy in modeling protein tertiary structures.[1,2] The precise estimation of inter-residue (atom) distances/ contacts in proteins constitutes a fundamental requirement for most protein structure prediction methodologies[3–5]

RaptorX is one of the early methods that achieved good accuracy in protein structure prediction by deep learning, which employed a ResNet architecture to predict all inter-residue contacts simultaneously.[6] Building on this success, RaptorX subsequently extended the approach to distance prediction which provides more comprehensive structural constraints than contact prediction alone.[7] Subsequently, distance prediction has been widely adopted for protein structure prediction, including in TrRosettaX,[8] ProFold,[9] DMPfold[10] and AlphaFold1[11] - the top-performing method in CASP13. These methods formulate distance prediction as a multi-class classification task, discretizing inter-residue distances into multiple bins. Alternatively, it was found that, with the same ResNet, real-valued prediction outperforms discrete-valued prediction in both contact and distance accuracy.[12] This finding has spurred growing interest in regression-based approaches, with methods like DeepDist and REALDIST demonstrating the potential of real-valued prediction distance prediction for high-accuracy structure modeling.[13,14]

Nevertheless, whether employing classification or regression frameworks, current distance prediction methods predominantly depend on coevolutionary information extracted from multiple sequence alignments (MSAs). This reliance stems from the well-established evolutionary principle that spatially proximal residues exhibit correlated mutation patterns, enabling the inference of inter-residue distance from MSAs.[15,16] While MSA-derived features (e.g., position-specific scoring matrices and covariance matrices) can be precomputed as model inputs, this conventional approach suffers from two key limitations: (1) incomplete extraction of evolutionarily informative patterns from MSAs, and (2) substantial computational demands.[17,18] A few publicly available methods attempt to effectively use the encoding of MSA as the direct input to model. For example, CopulaNet learns inter-residue distances through an MSA encoder directly;[9] RoseTTAFold and AlphaFold2 use direct embeddings of MSA to output atomic coordinates directly;[19,20] MSA Transformer is a protein language model that directly takes MSA as input, retaining the raw information of MSA while reducing computational costs.[21]

Even MSA-based strategy demonstrate strong predictive power for residue distances, it also suffers fundamental limitations that are particularly pronounced for proteins with poor sequence homology. The indirect correlations in MSA could lead to transitivity in residue spatial proximity and consequently cause incorrect estimation of inter-residue contacts/distances.[9] On the other side, the emergence of protein language models offers a new perspective—for example, ESMFold and OmegaFold extend these prediction techniques to even weakly homologous proteins by directly learning evolutionary information from sequences.[22–24] These methods eliminates the need for MSA and greatly simplified the neural architecture used for inference, showing great advantage to capture long-range interactions.

To effectively leverage the strengths of MSA derived coevolutionary features and sequence features by protein language model, this study developed MF-ProtDisMap (Multi-Feature Protein Distance Map), an integrative framework with multi-feature fusion for real-valued distance prediction. Specifically, MF-ProtDisMap extracts row and column attention features from the pretrained MSA Transformer to capture inter-residue interaction patterns and coevolutionary information, and incorporates ESM2 to learn protein sequence representations. Subsequently, group pooling is employed to reduce the dimensionality of the feature representations and fuse the two types of features. Inspired by the semantic modeling capability of diffusion models in self-supervised tasks — particularly their strength in capturing semantic information through implicit learning — we integrate diffusion-based modeling with a triangular attention mechanism to construct the Diff-former module.[25,26] This novel architecture significantly improved the representation of fused features. On the 4.05_release dataset (sourced from FreeProtMap),[27] MF-ProtDisMap achieves a Mean Absolute Error (MAE) of 2.20Å and a Root Mean Square Error (RMSE) of 3.40Å in real-valued protein distance prediction, outperforming ESMFold (which achieves an MAE of 2.56Å and an RMSE of 5.38Å). Ablation studies show that the integration of the Diff-former module improves the overall performance of MF-ProtDisMap. Moreover, MF-ProtDisMap outperforms models that rely solely on either MSA Transformer or ESM2 features, validating the effectiveness of our multi-feature fusion strategy. These findings suggest that a more comprehensive integration and utilization of sequence semantics with MSA-derived coevolutionary features leads to more accurate protein distance predictions.

## 2. Results

### 2.1. Overview of MF-ProtDisMap

We propose MF-ProtDisMap to enhance the accuracy of protein residue real-valued distance prediction by integrating protein sequence features with the coevolutionary features contained in MSA. The MF-ProtDisMap model consists of three main modules: the MSA feature generation module, the protein sequence feature generation module, and the Diff-former module, as shown in **Figure 1**.

**Figure 1.**
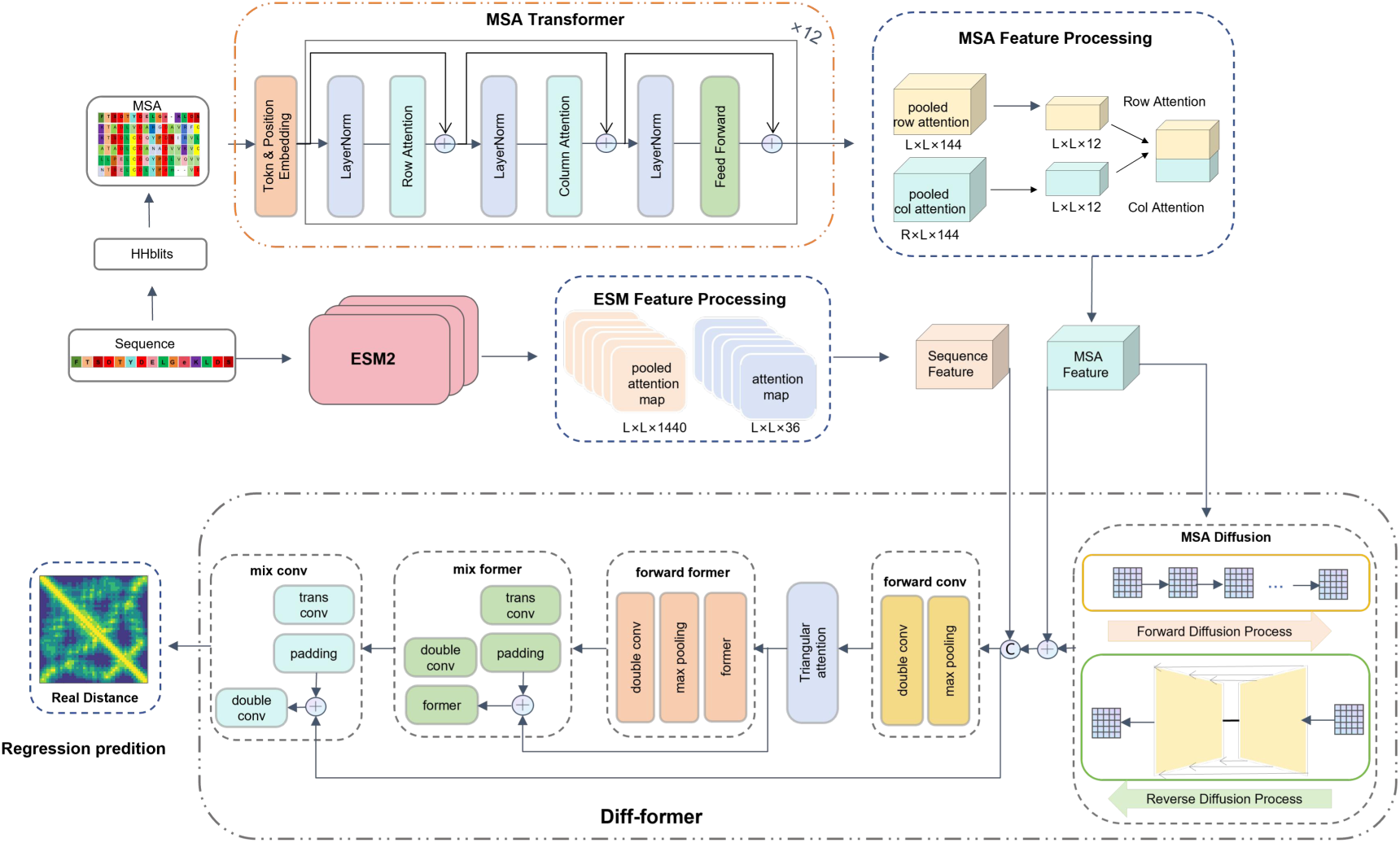
The MF-ProtDisMap Framework. MF-ProtDisMap consists of five main components: MSA Transformer, ESM2, MSA Feature Processing, ESM Feature Processing, and Diff-former. The MSA is input into the MSA Transformer, which extracts row attention and column attention features. These features are then processed through the MSA Feature Processing module, which includes dimensionality reduction and concatenation, to generate T-MSA features. The target sequence is fed into ESM2 to extract sequence-level attention features, which are processed through the ESM Feature Processing module via dimensionality reduction to obtain sequence features. The MSA Diffusion module further learns the T-MSA features to produce D-MSA features. Finally, the Diff-former module integrates the T-MSA features, D-MSA features, and sequence features to predict the real-valued inter-residue distances of the protein.

We employ HHblits to identify homologous sequences and construct MSA, which is subsequently processed by MSA Transformer to extract both row attention and column attention features, capturing residue-level coevolutionary relationships. Dimensionality reduction is then performed using group pooling, and the reduced features are projected into a unified 2D space and concatenated to form the T-MSA features, which emphasizes coevolutionary information between residues. In parallel, we extract sequence attention features using ESM2, and after processing with group pooling, these features—referred to as sequence features—primarily encode the semantic information of the target protein itself.

Diffusion models are not only used for data generation, the intermediate representations learned during the diffusion process can also benefit downstream prediction tasks.[28] Building on this insight, we designed the Diff-former module, which incorporates a triangular attention mechanism to capture the spatial invariance within the MSA and generate richer feature representations, referred to as D-MSA features. A residual connection is applied between the D-MSA and T-MSA features to implicitly align them within a unified feature space while preserving the original structural information from the MSA. These residual hybrid features are then concatenated with the sequence features obtained from ESM2, with the goal of maintaining feature diversity and enabling synergistic information enhancement. Finally, the concatenated features are fed into a Former network, which leverages a triangular attention mechanism to further learn a unified representation of geometric, evolutionary, and sequence information, thereby enabling accurate prediction of real-valued inter-residue distances.

### 2.2. Representation generation

#### 2.2.1. MSA features

To analyze a specific protein sequence, we employ HHblits (version3.3.0) in conjunction with the uniclust30_2020_03 database for generating MSA profiles for the sequences under investigation.[29,30] The resulting MSAs are used for subsequent feature extraction and analysis. Specifically, each MSA is represented as an *r×l* character matrix, where *r* denotes the number of homologous sequences and *l* represents the length of each sequence. To simplify the computation and reduce resource consumption, *r* is fixed at 64. This *r×l* matrix is subsequently input into the MSA Transformer for feature extraction. The MSA Transformer consists of 12 stacked encoder layers, each comprising 12 attention heads. This architecture enables the model to comprehensively capture coevolutionary information across sequences. Each attention block sequentially includes a row attention layer, a column attention layer, and a feed-forward layer, with each layer followed by layer normalization to ensure numerical stability.

We extract two types of features—row attention and column attention—from the attention modules of the MSA Transformer and construct a two-dimensional feature matrix. Specifically, we obtain a row attention weight matrix of shape *L×L×144* from the final attention layer of the MSA Transformer. To reduce noise and preserve key information, we apply group pooling to perform dimensionality reduction, resulting in a feature matrix of shape *L×L×12*. Meanwhile, a column attention weight matrix of shape *R×L×144* is also extracted from the final attention layer. This matrix is subjected to dimensionality reduction and subsequently mapped to a tensor of shape *L×L×12* to optimize feature representation. Finally, the row-attention features and column-attention features are concatenated along the feature dimensions to form a two-dimensional feature map of *L×L×24*, referred to as the T-MSA features. Chen et al. processed the features extracted from the MSA Transformer into two feature types: MSA Features and Row Attention Maps.[31–33] To evaluate the effectiveness of different feature processing methods, this study conducts comparative experiments, and the results show that our method performs better.

#### 2.2.2. Sequence features

In this study, we input protein sequences of length *l* into ESM2 for feature encoding. ESM2 is built on the Transformer architecture, comprising multiple self-attention layers and feed-forward neural networks. Through pretraining, ESM2 effectively captures both the evolutionary conservation and variability within protein sequences. Moreover, the self-attention mechanism of ESM2 allows it to learn long-range dependencies within the sequence, extracting global information by calculating the relationships between positions across the entire sequence.

However, the high-dimensional sparse attention matrices generated by ESM2 significantly increase computational costs, while simple dimensionality reduction methods may result in the loss of a large amount of important information.Therefore, this study employs a group pooling approach to extract more information representations. Specifically, the sequence features generated by ESM2 are initially represented as a tensor of size *L×L×1440*, which is compressed into a feature representation of size *L×L×36* using group pooling. The advantage of the group pooling method lies in its ability to reduce the dimensionality of attention maps within each feature subspace individually, thereby minimizing redundancy while maximizing the retention of valuable information.

### 2.3. Diff-former module

Based on the triangular attention constraint, we design a hybrid structure called Diff-former. Considering the advantages of diffusion models in unsupervised semantic representation learning—specifically, their ability to learn meaningful data representations that enhance downstream prediction tasks—we feed the 2D feature map XT, constructed by MSA Transformer, into the diffusion module Diff(.). Here,Xt denotes the intermediate state of XT at time step t (0 ≤ t ≤ T). Two Markov processes are defined in the module: the forward diffusion process gradually adds noise to X0, while a the reverse generation process iteratively samples from the prior distribution XT. We utilize the denoising diffusion probability model (DDPM) for generation.[34] Conditioning on an initial distribution, where X is a continuous variable, the forward process is defined by gradually diffusing X0 to a normal prior p(LT ) ∼ N (0, I) via q(Xt|XtX ):

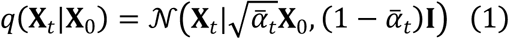

The backward generation process is given by the following equation:

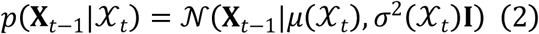

Where β denotes the noise variance series, 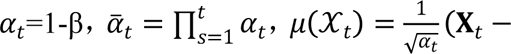 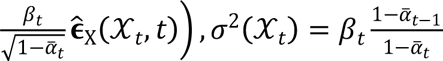 Denoising term 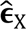 is predicted using the UNet model. Through the noise-adding and denoising process of the diffusion model, the representational capacity of the MSA Features is enhanced, enabling more effective capture of underlying patterns and trends of protein interactions. After generating the features XD from the Diff module, we apply a residual connection to combine them with the MSA Transformer features XT, resulting in an enhanced feature representation XTD. This process not only preserves the MSA structural information but also implicitly aligns the D-MSA features generated by the Diff module into the MSA feature space, thereby improving the model’s feature representation ability.

The basic architecture of Former consists of a bottleneck structure and residual structures. The bottleneck structure is designed to capture robust and high-dimensional representations of the input while minimizing the risk of overfitting.[27] The residual structure facilitates easier learning by allowing each neural network block to focus on modeling only the residuals between the input and output.

The triangular attention module is set on the second layer of the Former to balance computational costs and prediction accuracy. Here, XE represents the sequence features extracted from ESM2. The module forward_conv refers to the forward CNN, which consists of a max-pooling layer followed by a two-layer CNN. The mix_conv module is composed of a transposed CNN, padding, and a double CNN. forward_former and mix_former add the former module on top of forward_conv and mix_conv respectively. The detailed process is shown in **Equation (3)**. We fuse the residual-connected features XTD with the sequence features XE through feature concatenation, aligning features from different sources within a unified space, and feed the result into the Former module. By leveraging a triangular attention mechanism, the Former module learns a joint representation of geometric, evolutionary, and sequential information, thereby enabling accurate prediction of protein inter-residue distances.

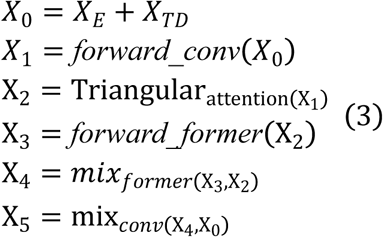

The window size of the maximum pooling layer is set to 2, and the number of filters in forward_conv, forward_former, mix_former, and mix_conv are set to (64,64), (128,128), (64,64), and (36,36), respectively.

### 2.4. Evaluation of Real-Valued Distance Prediction Performance

#### 2.4.1. Comparison with other methods

To comprehensively evaluate the performance of MF-ProtDisMap in the task of protein real-valued distance prediction, we conducted a comparative analysis against state-of-the-art structure prediction methods.[35] As the latest open-source models are primarily designed for protein structure prediction rather than directly outputting real-valued distances, we converted their predicted 3D structures into distance to ensure objectivity and consistency in the comparative experiments. For evaluation, we employed widely used regression metrics, including Mean Absolute Error (MAE), Root Mean Square Error (RMSE), and Coefficient of Determination (R²). In addition, we calculated their mean deviations (S_MAE, S_RMSE, and S_R2), to assess the stability of the model. The experimental results are presented in Figure 2, with the best performance for each metric highlighted in bold.

**Figure 2.**
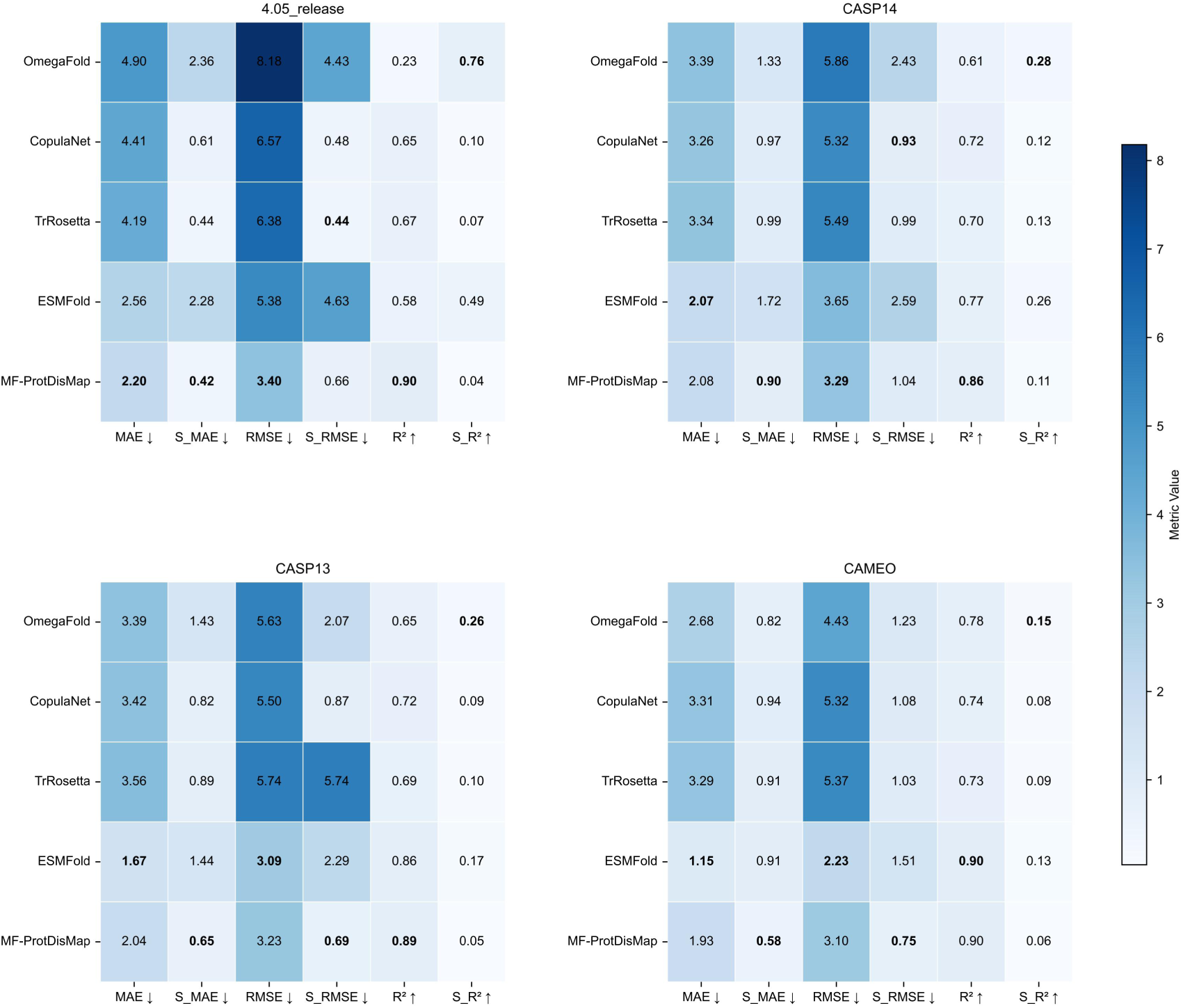
Comparative evaluation of real-valued distance prediction methods on four benchmark datasets. The x-axis represents evaluation metrics, including MAE, S_MAE, RMSE, S_RMSE, R², and S_R², while the y-axis lists the model names. Subfigures correspond to the 4.05_release, CASP14, CASP13, and CAMEO datasets, respectively. Color intensity indicates metric values, with lighter colors representing smaller values. The best-performing values are highlighted in bold. For all metrics except R², smaller values are better; for R², larger values indicate better performance.

As shown in Figure 2, MF-ProtDisMap achieves a MAE of 2.20 Å, an RMSE of 3.40 Å, and an R² of 0.90 on the 4.05_release dataset, indicating a low prediction error. Furthermore, the mean deviations of each metric demonstrate the model’s robustness in handling new proteins. Specifically, MF-ProtDisMap achieved an S_MAE of 0.42 Å, an S_RMSE of 0.66 Å, and an S_R2 of 0.04 Å on the 4.05_release dataset. We further compared MF-ProtDisMap with other state-of-the-art methods on the CASP14, CASP13, and CAMEO datasets. On the CASP13 dataset, MF-ProtDisMap achieved an S_MAE of 0.65 Å, an S_RMSE of 0.69 Å, and an S_R² of 0.05, outperforming the best results (S_MAE = 0.82, S_RMSE = 0.87, S_R² = 0.09). On the CAMEO dataset, MF-ProtDisMap also demonstrated superior performance with an S_MAE of 0.58 Å, an S_RMSE of 0.75 Å, and an S_R² of 0.06, exceeding the best results (S_MAE = 0.82, S_RMSE = 1.03, S_R² = 0.08). These results strongly validate the effectiveness and advantages of MF-ProtDisMap in the task of real-valued distance prediction.

#### 2.4.2. Error Evaluation of MF-ProtDisMap in Real-Valued Distance Prediction

From the perspective of structure and binding site prediction, inter-residue interactions carry greater significance than non-interacting pairs, with short-range interactions and long-range ones.[15] To evaluate the model’s performance in predicting real-valued distances between residue pairs, we categorized the residue pairs into three groups based on their true distances: short-range (<6 Å), medium-range (6-12 Å), and long-range (≥12 Å). We then computed the MAE for each group on the 4.05_release dataset. **Figure 3(a)** displays the MAE distribution through boxplots across different distance ranges. The model exhibits optimal performance for short-range interactions (median error = 2.23 Å). The error slightly increases for medium-range interactions (median error = 3.08 Å) and is highest for long-range interactions (median error = 6.01 Å), demonstrating robust capability in capturing both local and global structural features.

**Figure 3.**
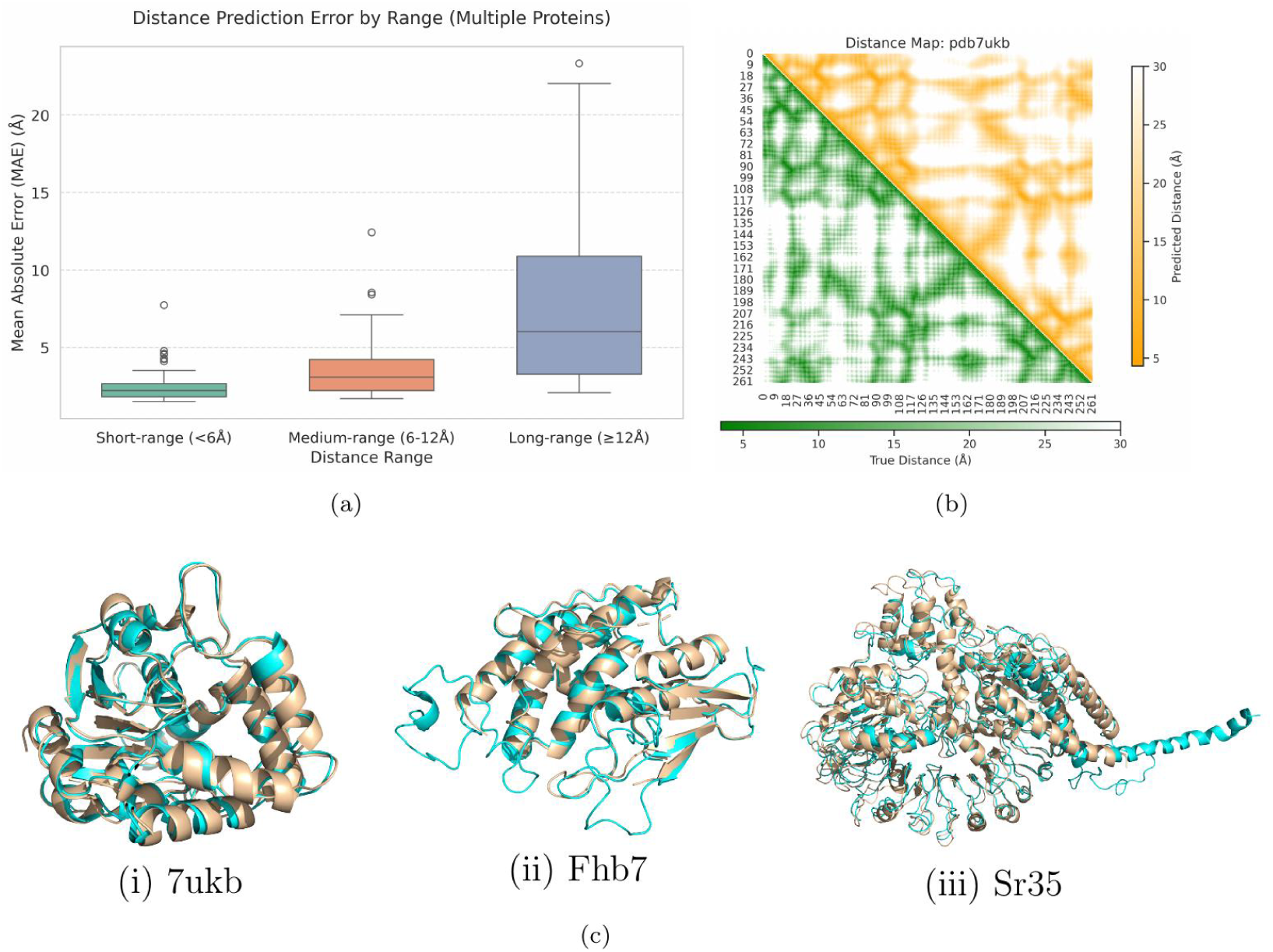
Performance of MF-ProtDisMap in Protein Real-Valued Distance Prediction. (a) Box plot of the MAE distribution across different distance ranges in the test set, where residue pairs are categorized into short-range (<6 Å), medium-range (6–12 Å), and long-range (≥12 Å) based on their true distances. (b) Heatmap visualization of the true distance matrix (lower-left) and the predicted distance matrix (upper-right) for target T1024. (c) Representative 3D structural models of (i) ancestral plant α/β-hydrolase (PDB: 7UKB), (ii) the major wheat scab resistance protein Fhb7, and (iii) the wheat stem rust resistance receptor Sr35, reconstructed by ProtDisFold using real-valued inter-residue distance constraints.

Given the lack of available tools capable of reconstructing protein structures directly from real-valued distance predictions, we developed a preliminary modeling tool based on PyRosetta for distance-constrained structure modeling. Since the current framework does not incorporate complete backbone conformational constraints (such as dihedral angles), we extracted φ/ψ dihedral angle information from native PDB structures to assist in structure reconstruction. The tertiary structures of example protein 7ukb, Fhb7-GST and a plant NLR Sr35, with accurate prediction in distance matrix, were successfully reconstructed (**Figure 3b, c)**, demonstrating that accurate real-valued distance predictions can be used as constraints for protein structure modeling in the future.

### 2.5. Evaluation of Contact Prediction Performance

To further evaluate the performance of MF-ProtDisMap, we converted the generated distance maps into threshold-based contact maps and conducted a comparative analysis with existing residue contact prediction methods (ESM-1b, SPOT-Contact-LM),[36,37] protein language models (OmegaFold),[23] and CopulaNet,[9] which uses full MSA as input. The experimental results are summarized in **Table 1**. On the 4.05_release dataset, MF-ProtDisMap achieved an ROC value of 84.56%, PR of 81.01%, F1 of 72.86%, Precision of 86.25%, and Recall of 63.28%. Compared to the second-best results (PR of 72.25%, F1 of 64.76%, Precision of 83.44%, Recall of 59.74%), MF-ProtDisMap shows significant improvement. To further assess its performance, we evaluated MF-ProtDisMap on the CASP14, CASP13, and CAMEO datasets. As shown in **Figure 4**, MF-ProtDisMap outperformed other models in PR and Precision metrics.

**Figure 4.**
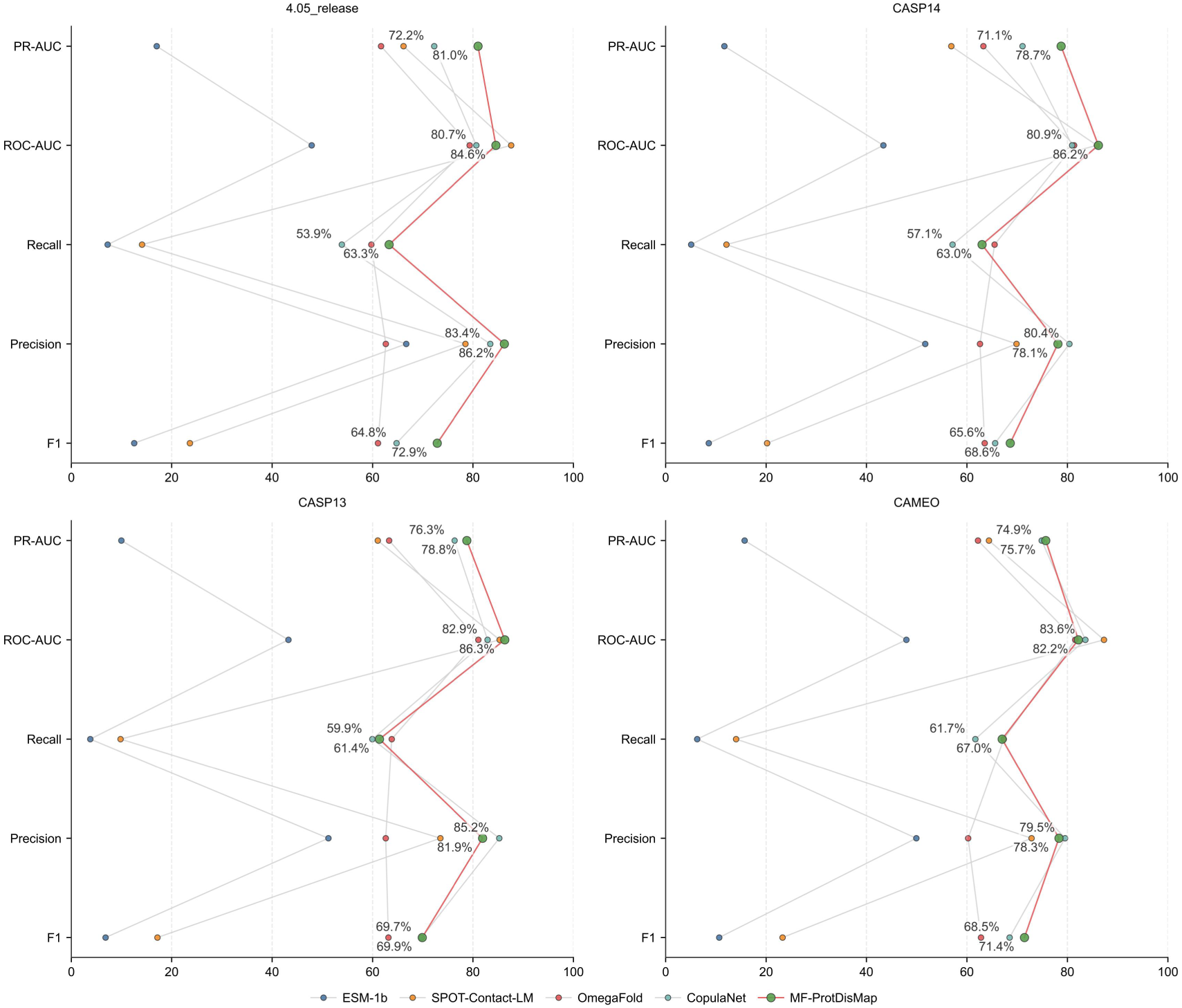
Comparative evaluation of inter-residue contact prediction methods on four benchmark datasets. The subfigures correspond to the 4.05_release, CASP14, CASP13, and CAMEO datasets. The x-axis represents the performance values of the models, in percentage (%), while the y-axis represents the different evaluation metrics. Different colored lines and points represent different models, with the results of MF-ProtDisMap highlighted in red.

**Table 1.**
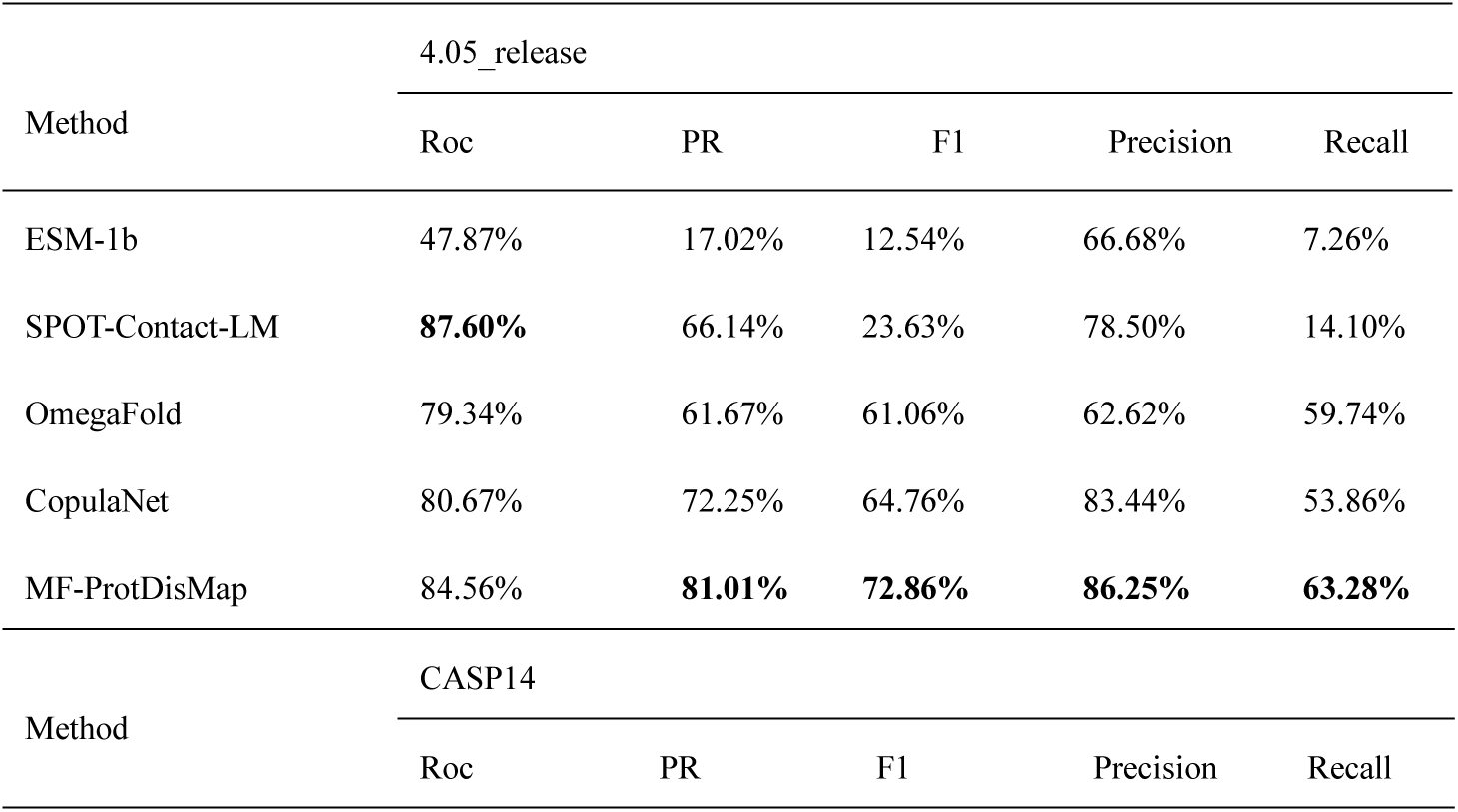
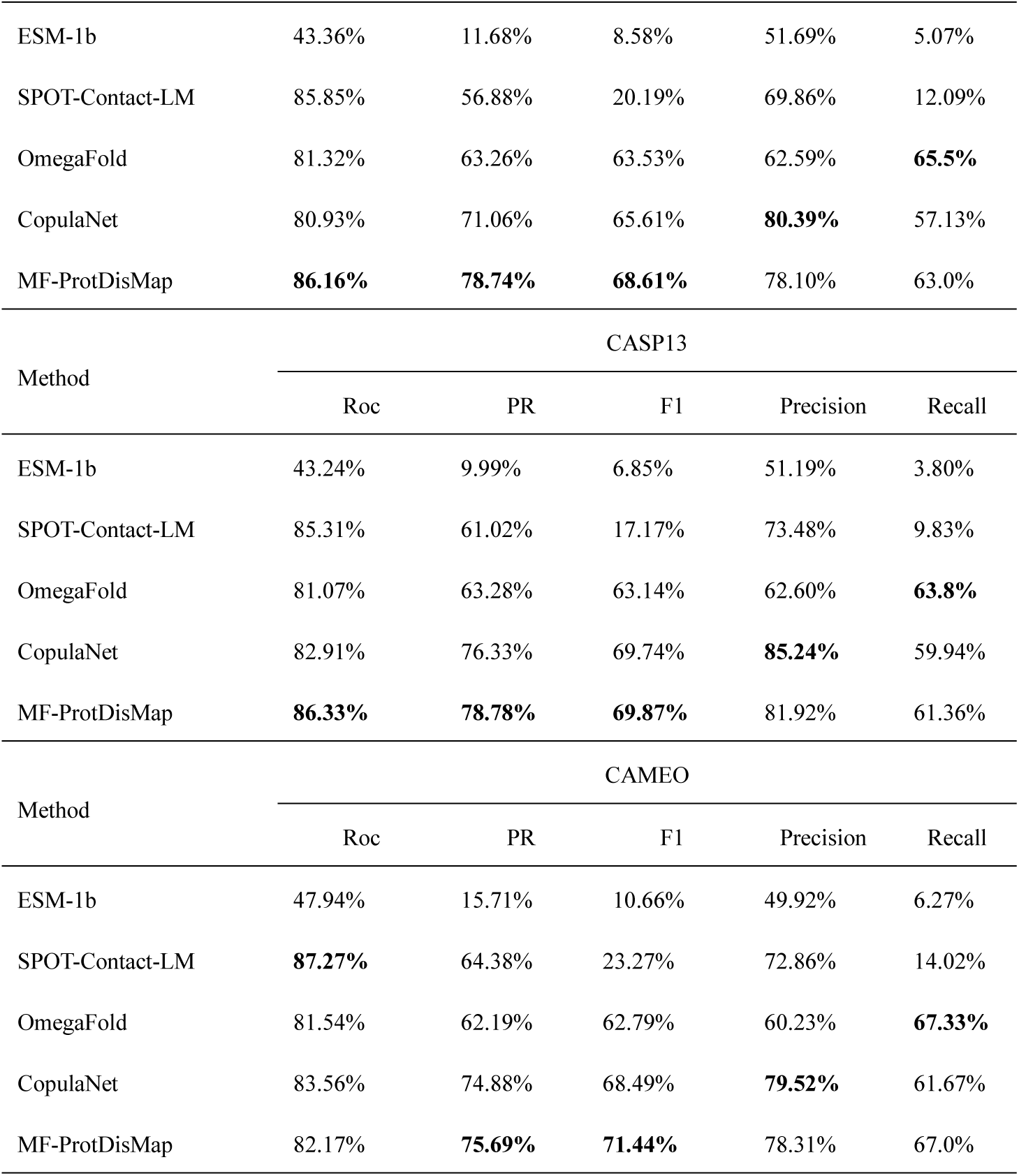
Evaluation results of MF-ProtDisMap with other models on 4.05_release, CASP13, CASP14, CAMEO datasets for classification tasks.

### 2.6. Ablation Experiments

In this study, we conducted a series of ablation experiments to evaluate the contributions of different input features and module designs to the overall performance of our model. By progressively adding or replacing key components within the architecture, we analyzed their individual impacts on both regression and classification tasks, thereby validating their necessity and effectiveness. Specifically, when using only the features extracted by the MSA Transformer, the model demonstrated relatively poor performance on the 4.05_release dataset, with a mean absolute error (MAE) of 2.47Å and a root mean square error (RMSE) of 3.81Å in the regression task, and classification metrics of ROC = 69.90%, PR = 68.47%, F1 = 52.82%, and Recall = 38.75%. When using only the sequence features extracted by ESM2, the model achieved moderate improvements, with MAE and RMSE reduced to 2.43Å and 3.73Å, respectively. The ROC increased to 81.26%, while the precision-recall area (PR) reached 77.52%, with F1 and Recall improving to 63.63% and 52.02%, respectively. (**Table 2** and **Table 3**)

**Table 2.**
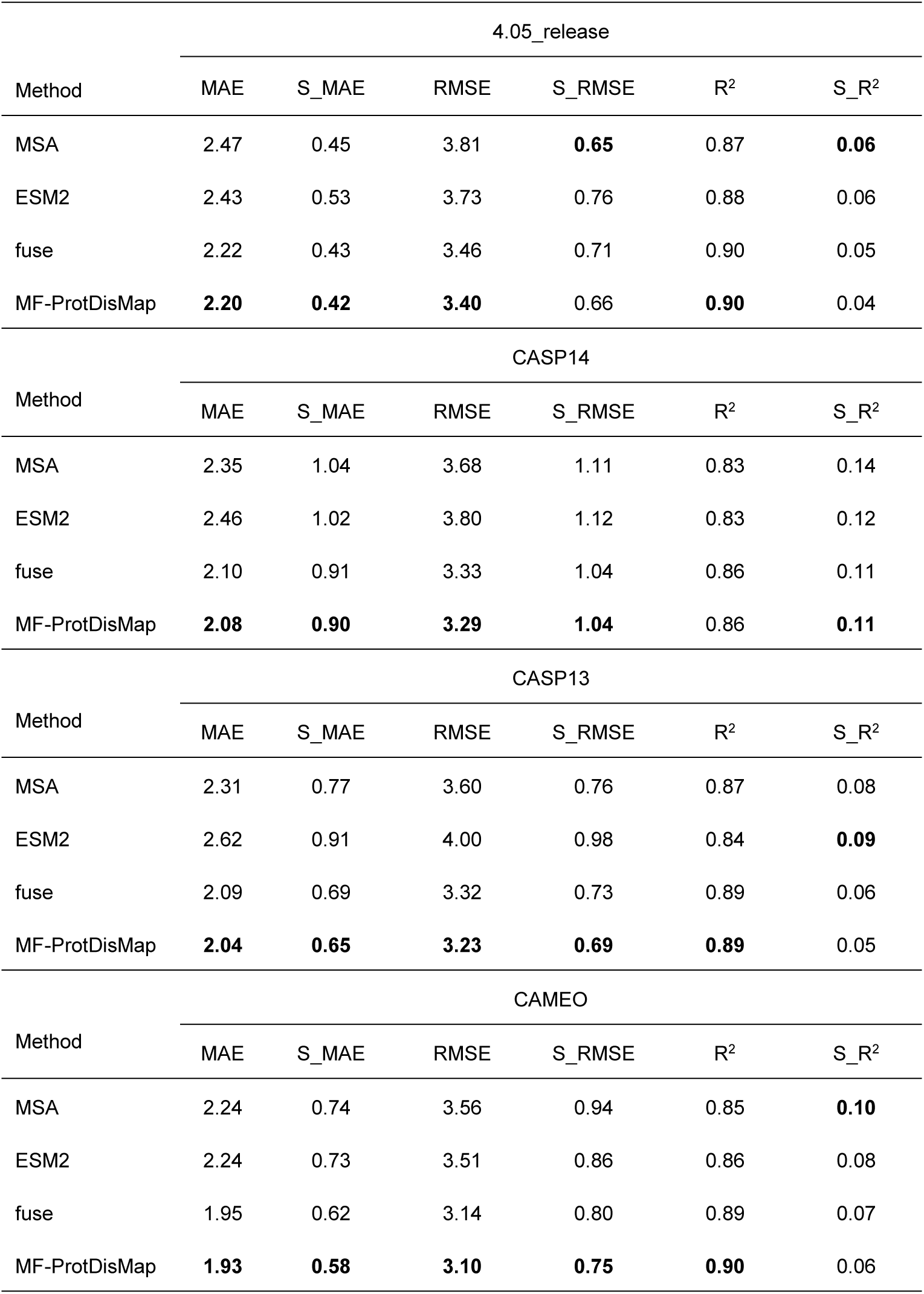
Evaluation results of regression metrics for ablation experiments.

**Table 3.** Results of the assessment of classification indicators for ablation experiments.

| Method | 4.05_release |  |  |  |  |
| --- | --- | --- | --- | --- | --- |
|  | Roc | PR | F1 | Precision | Recall |
| MSA | 69.90% | 68.47% | 52.82% | **89.12%** | 38.75% |
| ESM | 81.26% | 77.52% | 63.63% | 84.37% | 52.02% |
| fuse | 82.75% | 78.76% | 70.56% | 84.87% | 60.95% |
| MF-ProtDisMap | **84.56%** | **81.01%** | **72.86%** | 86.25% | **63.28%** |

| CASP14 |  |  |  |  |  |
| --- | --- | --- | --- | --- | --- |
| Method | Roc | PR | F1 | Precision | Recall |
| MSA | 69.68% | 63.16% | 45.16% | **83.33%** | 33.85% |
| ESM | 80.59% | 74.10% | 59.76% | 76.78% | 51.47% |
| fuse | 84.22% | 75.75% | 65.47% | 78.07% | 58.67% |
| MF-ProtDisMap | **86.16%** | **78.74%** | **68.61%** | 78.10% | **63.06%** |

| CASP13 |  |  |  |  |  |
| --- | --- | --- | --- | --- | --- |
| Method | Roc | PR | F1 | Precision | Recall |
| MSA | 62.20% | 57.14% | 42.97% | **86.53%** | 29.94% |
| ESM | 78.52% | 70.39% | 58.26% | 78.47% | 47.71% |
| fuse | 84.99% | 76.85% | 64.31% | 81.94% | 54.9% |
| MF-ProtDisMap | **86.33%** | **78.78%** | **69.87%** | 81.92% | **61.36%** |

| CAMEO |  |  |  |  |  |
| --- | --- | --- | --- | --- | --- |
| Method | Roc | PR | F1 | Precision | Recall |
| MSA | 58.40% | 58.47% | 47.79% | **83.44%** | 35.45% |
| ESM | 80.82% | 74.44% | 62.15% | 78.19% | 53.36% |
| fuse | 80.97% | 74.36% | 67.96% | 78.97% | 61.37% |
| MF-ProtDisMap | **82.17%** | **75.69%** | **71.44%** | 78.31% | **67.06%** |

Further improvements were observed when combining MSA and ESM2 features (denoted as “fuse” in Tables 3 and 4). In this setting, the model achieved lower MAE and RMSE values of 2.22Å and 3.46Å, respectively, while F1 and Recall increased to 70.56% and 60.95%. (**Table 2** and **Table 3**) These results indicate that the integration of coevolutionary features from MSA and semantic information from ESM2 provides complementary benefits, significantly enhancing the model’s prediction capability.

Finally, building upon the aforementioned configuration, we introduced the Diff-former module to construct the full MF-ProtDisMap model, which led to further performance improvements. On the 4.05_release dataset, MF-ProtDisMap achieved an MAE of 2.20Å and an RMSE of 3.40Å in the regression task. For the classification task, the model reached a ROC of 84.56%, a PR of 81.01%, an F1 score of 72.86%, and a Recall of 63.28%. Compared to the “fuse” setting, MF-ProtDisMap reduced MAE and RMSE by 0.02Å and 0.06Å, respectively, while improving F1 and Recall by 2.30% and 2.33%. (**Table 2** and **Table 3**)These results demonstrate that the Diffusion module, through deep modeling of the feature space, enables the learning of more meaningful and informative representations. Consequently, it enhances both the prediction accuracy and the generalization capability of the model.

Moreover, to evaluate the generalizability of the proposed module design and feature combination strategy, we conducted ablation experiments on three additional public benchmark datasets: CASP14, CASP13, and CAMEO. The results on these datasets were consistent with those observed on the 4.05_release dataset, further validating the importance of each model component and the rationality of the overall architectural design. Specifically, the model performed worse when using only MSA or ESM2 features individually, compared to the fused feature setting. In contrast, the incorporation of the Diff-former module consistently enhanced performance across all datasets, with particularly notable improvements in classification metrics such as F1 score and Recall. These findings demonstrate that MF-ProtDisMap possesses strong adaptability and robustness, enabling accurate real-valued distance and contact prediction across diverse data distributions.

## 3. Discussion

The accuracy of protein structure prediction has always been a central issue in the field of computational biology, and significant progress has been made in recent years with the advancement of deep learning technologies.[38–40] However, the limited performance of protein distance prediction still constrains the final performance of structural prediction tools. Compared to inter-residue distance prediction, which is of a categorical nature, the research on real-valued distance prediction is relatively limited, but it holds great potential in capturing more refined interactions between protein residues.[41,42] Here, our MF-ProtDisMap by merging language model and diffusion model succefully acchieved both accurate real-valued distance prediction and contact prediction, offering an interpretable, adjustable, and scalable intermediate representation for structure prediction.

The use of MSA-derived coevolutionary features has significantly advanced protein structure prediction, demonstrating particular efficacy for proximal residue distance estimation. However, this approach shows substantially reduced effectiveness for proteins with poor sequence homology.[43,44] On the other hand, recent application of language models in protein structure prediction like ESM2 demonstrates powerful sequence semantic capture abilities, especially in the prediction of diverged sequences and long-range interactions.[45–48] However, the large amount of sequence semantic information may mask the coevolutionary features and spatial invariance that was advanced in MSA. It is proposed that a multi-feature model by integrating the sequence semantic features and coevolutionary information from MSA can effectively leverage the strengths of both methods. This is underpined by the result of ablation experiments that the fused model achieves an MAE of 2.22 Å and an RMSE of 3.46 Å in the regression task, better than either MSA or the sequence language model alone (Table 2). It thus indicate a new perspective for designing more efficient protein representation learning methods with fusion strategy.

As protein real-valued distance prediction model, MF-ProtDisMap not only exhibit advantage with fused strength of MSA and sequence semantic features, but also demonstrates significant computational resource advantages, requiring only a single A100 GPU for training, greatly reducing computational costs. Additionally, the model’s performance on multiple public datasets, including CASP13, CASP14, and CAMEO, demonstrates its strong generalization capability, confirming its stability and effectiveness across diverse scenarios. This can be attributed to the following key factors: (1) By utilizing the noise addition and denoising process of the diffusion model in self-supervised tasks, an implicit representation space is constructed. During the backward denoising process, the network dynamically reconstructs the data distribution, effectively capturing the common semantic features of the MSA. (2) MF-ProtDisMap incorporates the triangular attention module from AlphaFold-2, which effectively captures the geometric relationships between residues in the distance map; (3) Feature dimensionality is reduced using group pooling, which decreases the dimensionality of attention maps in each feature subspace separately, ensuring maximal information retention while reducing redundancy.

In summary, the multi-feature model of MF-ProtDisMap significantly improves the accuracy of protein distance prediction when using standard database, that paves a way for the development of efficient and computationally lightweight deep learning models. While the current dataset is sufficient for feature extraction and model training, its relatively limited size can constrain the model’s generalization ability and accuracy. Future research should employ expanded datasets and integrate more comprehensive biological knowledge. A more deep exploration of MSA’s coevolutionary features and spatial invariance, along with integration with more comprehensive large language models, should favorate to develop more efficient protein representation learning models.

## 4. Materials and Methods

### 4.1. Dataset and MSA Generation

We employed the dataset curated by Yang et al. as the training dataset, which originally consists of 15051 protein sequences.[35] To satisfy the input constraints of the ESM2 language model, sequences exceeding 1,022 residues were excluded, resulting in a final dataset of 14,942 sequences. For model training, 11,000 sequences were randomly selected as the training set, with the remaining 3,942 sequences used for validation. The model was implemented using the PyTorch framework and trained on an Nvidia H800 GPU. An initial learning rate of 1e^-3^ and a batch size of 1 were used. Parameter optimization was performed using the Adam optimizer with a weight decay of 0.01.

To evaluate the performance of the model on newly discovered proteins, we employed the dataset 4.05_release compiled by Tang et al. as one of the test sets, which contains 90 proteins.[27] In addition, to assess the model’s generalization capability across different datasets, we further utilized benchmark test sets provided by the CASP13, CASP14, and CAMEO competitions. Since a complete protein dataset was not available for the CASP15 competition, we did not report any related testing results. Specifically, our CASP13 dataset includes 40 proteins from the CASP13 competition, CASP14 comprises 138 proteins from CASP14, and CAMEO contains 129 proteins released by the CAMEO competition (June to September 2020). In these datasets, some proteins have incomplete structural information, particularly in CASP13, CASP14, and CAMEO.

To generate the MSA for a query protein sequence, we utilized the HHblits tool (version 3.3.0) to search against the UniRef30_2020_03 database. The resulting MSAs were saved in A3M format and subsequently used as input for the MSA Transformer.

### 4.2. Performance Metrics

For the evaluation of real-valued distance prediction, we adopted commonly used regression metrics, including Mean Absolute Error (MAE), Root Mean Square Error (RMSE), and the Coefficient of Determination(R-Square,R^2^). The corresponding evaluation formulas (**Equations 4–7**) are provided below, where *d* denotes the ground truth inter-residue distance, *d̂* represents the predicted inter-residue distance, *d̄* is the mean of the true distances, and *s* represents the statistics of MAE, RMSE, and R^2^.

For the evaluation of contact prediction, we adopted the standard CASP definition, where two residues are considered to be in contact if the distance between their C*β* atomic is less than 8.0 Å. To further assess the performance of MF-ProtDisMap, we converted the predicted distance matrices into binary contact matrices based on this threshold and compared MF-ProtDisMap with other state-of-the-art residue contact prediction methods. During the evaluation process, we employed commonly used metrics for classification tasks, including ROC-AUC (Area Under the ROC Curve), PR-AUC (Area Under the Precision-Recall Curve), Precision, Recall, and F1 score, as detailed in **Equations 8-10**. Moreover, the SmoothL1 Loss function is a smoothed L1 loss function that combines the advantages of squared and absolute losses, effectively reduces the instability of the values as they approach zero. Its mathematical formulation is given below (**Equation 11**).

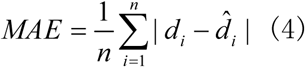

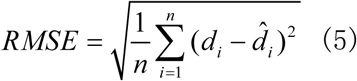

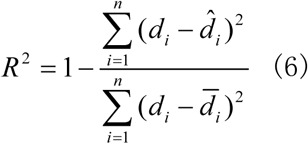

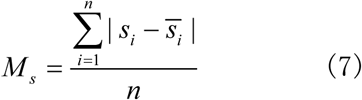

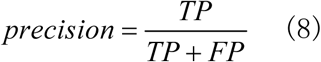

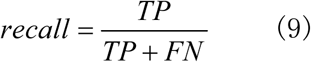

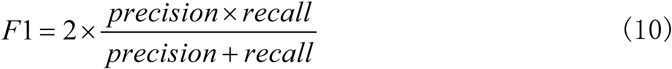

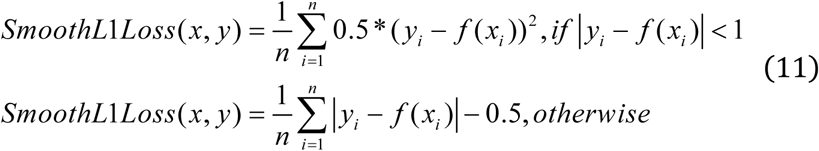

### 4.3. Subsampling of MSA

During the training phase, we implemented a subsampling method to process MSAs, with the dual purpose of enhancing model robustness through data augmentation and mitigating computational burden to prevent GPU memory overload caused by large-scale alignments. Specifically, instead of feeding the complete MSA directly into the model during training, we selected a subset of the MSA to ensure computational efficiency and alignment with the model’s input constraints. This subsampling approach randomly selects a number of sequences from the MSA, retaining at least 16 rows, with the query sequence consistently placed in the first row.

During the inference phase, we likewise refrain from inputting the full MSA into the model, primarily due to its high computational cost and the potential for performance degradation when the input size significantly exceeds that used during training. To address this, we explored three subsampling strategies: random selection, diversity maximization, and diversity minimization. Among these, the diversity-maximizing strategy achieved the most favorable balance between computational efficiency and the representativeness of the selected sequences.

Specifically, this approach calculates the average Hamming distance for each sequence and prioritizes those with higher distances, thereby enhancing the diversity of the subset. Ultimately, a subset of 64 sequences is selected from the original MSA and used as the model input during inference.

### 4.4. Tertiary structure remodeling

Given the current lack of tools that can directly reconstruct protein structures based on real distance prediction results, we have tentatively developed ProtDisFold— a PyRosetta-based tool for distance constraint-driven protein structure modeling.[49] Specifically, the predicted real-valued distances are converted into distance restraints required by Rosetta,[50] with upper and lower bounds set to 15.0 Å and 2.0 Å, respectively, and are subsequently used for 3D structure modeling under the Rosetta ab initio protocol. In addition, the tool can leverage dihedral angle information from known structures and convert it into a von Mises distribution, providing additional prior knowledge for subsequent conformational sampling.

ProtDisFold first performs probabilistic sampling of the backbone dihedral angles and generates an initial conformation based on distance constraints. Conformational optimization is then carried out using a two-stage strategy: The first stage involves coarse optimization in centroid mode. A progressive angular perturbation strategy is employed, with the perturbation range gradually decreasing from 20° to 4°. After each perturbation, energy minimization is performed using the LBFGS algorithm, and the process moves to the next stage once the energy score stabilizes. The second stage switches to full-atom mode for refinement, allowing full freedom of adjustment for both the backbone and side chains, and performs multiple iterations of optimization using the FastRelax protocol.

To improve modeling efficiency, we adopted a parallel modeling mechanism, generating multiple candidate structures simultaneously and performing random initialization for each model to explore the conformational space. In the model evaluation phase, ProtDisFold uses the TMscore program to assess the quality of all candidate structures relative to the reference structure, calculates the TM-score, and ranks the models. The highest-scoring model is selected as the final prediction result. The entire process automatically records the scores of each model and generates an evaluation report.

## Data Availability Statement

The training dataset used in this study is based on the benchmark dataset released by Yang et al., while the testing datasets were obtained from the dataset released by Tang et al., as well as public datasets from CASP and CAMEO. All original datasets are available through the corresponding published papers or official websites.

## Acknowledgements

This work was supported by the National Key Research and Development Program of China (2022YFF1001504), the National Natural Science Foundation of China (U23A20181), the National Natural Science Foundation of China (No. 62202281).

